# EXPLORING THE EFFECTS OF METABOLISM-DISRUPTING CHEMICALS ON PANCREATIC α-CELL BIOLOGY: A SCREENING TESTING APPROACH

**DOI:** 10.1101/2022.11.07.515444

**Authors:** Ruba Al-Abdulla, Hilda Ferrero, Talía Boronat-Belda, Sergi Soriano, Iván Quesada, Paloma Alonso-Magdalena

**Affiliations:** Instituto de Investigación, Desarrollo e Innovación en Biotecnología Sanitaria de Elche (IDiBE), Universidad Miguel Hernández, Elche, Spain; Centro de Investigación Biomédica en Red de Diabetes y Enfermedades Metabólicas Asociadas (CIBERDEM), Spain; Departamento de Fisiología, Genética y Microbiología, Universidad de Alicante, Alicante, Spain

**Keywords:** metabolism-disrupting chemicals, pancreatic α-cell, diabetes, metabolic diseases, screening testing

## Abstract

Humans are constantly exposed to many environmental pollutants, some of which have been largely acknowledged as key factors in the development of metabolic disorders such as diabetes and obesity. These chemicals have been classified as endocrine-disrupting chemicals (EDCs) and, more recently, since they can interfere with metabolic functions, they have been renamed as metabolism-disrupting chemicals (MDCs). MDCs are present in many consumer products, including food packaging, personal care products, plastic bottles and containers, and detergents. The scientific literature has ever-increasingly focused on insulin-releasing pancreatic β-cells as one of the main targets for MDCs. Evidence highlights that these substances may disrupt glucose homeostasis, altering pancreatic β-cell physiology. However, their potential impact on glucagon-secreting pancreatic α-cells remains poorly known despite the essential role that this cellular type plays in controlling glucose metabolism. In the present study, we have selected seven paradigmatic EDCs representing major toxic classes, including bisphenols, phthalates, perfluorinated compounds, metals, and pesticides. By using an in vitro cell-based model, the pancreatic α-cell line αTC1-9, we have explored the effects of these compounds on pancreatic α-cell viability, gene expression, and secretion. Our results indicated that most of the selected chemicals studied caused functional alterations in pancreatic α-cells. Moreover, we revealed, for the first time, their direct effects on key molecular aspects of pancreatic α-cell biology.

## 1. Introduction

Pancreatic α and β-cells are the most abundant endocrine cells within the islets of Langerhans, and together, they orchestrate a bi-hormonal secreting system central to the regulation of glucose metabolism. Specifically, pancreatic β-cells secrete insulin in response to elevated blood glucose levels, while α-cells release glucagon as a counter-regulatory hormone in response to hypoglycaemia conditions [1, 2]. Indeed, the adequate function of pancreatic α-cells and glucagon release constitute the first line of defence against hypoglycaemia. Beyond their function as glucagon-secreting cells, it is also known that pancreatic α-cells may behave as important keepers of pancreatic β-cells. For instance, in conditions of extensive β-cell loss, α-cells may convert into new functional β-cells, a process called transdifferentiation [3, 4]. Pancreatic α-cells may also exhibit a significant degree of plasticity in the form of an increased rate of proliferation to compensate when glucagon signalling is impaired [5-8]. Furthermore, hyperplastic α-cells seem to be able to overexpress the glucagon-like peptide GLP-1 [9], which is known to increase pancreatic β-cell proliferation and insulin secretion [10], thus contributing to replenishing insulin-producing cells. In addition to GLP-1, glucagon, as well as other paracrine signals from the pancreatic α-cell, play a beneficial role in regulating β-cell function [11, 12].

Impaired α-cell function and glucagon release are involved in the aetiology of diabetes [13]. In type 1 (T1D) and advanced type 2 diabetes (T2D), pancreatic α-cells fail to respond adequately to decreased plasma glucose levels, increasing the risk of hypoglycaemia [13]. On the contrary, in the context of β-cell failure, impaired inhibition of α-cell secretion at high glucose levels can contribute to postprandial hyperglucagonemia and hyperglycaemia in diabetes [13]. Overall, this evidence supports the critical importance of pancreatic α-cells not only in the maintenance of metabolic homeostasis but also in the pathophysiology of diabetes.

Diabetes is a complex disease caused by a combination of genetic and environmental factors. Among those environmental factors, a subset of chemicals named metabolism-disrupting chemicals (MDCs) has emerged as key players in the occurrence of this disease, particularly for T2D [14-17]. MDCs are found in many everyday products, including plastic bottles and food containers, food-can liners, cosmetics, flame retardants, and pesticides [14]. In recent years, it has been acknowledged that some of these chemicals can induce pancreatic β-cell malfunction and impair insulin action, thus playing a causative role in metabolic disorders progression [15-18]. While pancreatic β-cells are gaining importance as a main target of MDCs’ action, the potential impact of MDCs on pancreatic α-cell biology remains largely unknown. Besides this knowledge gap, another major limitation is that there are no effective methods with which to test for MDCs, and the regulatory tests currently employed in the European Union do not assess endocrine pathways related to the pancreatic islet system [19].

In the present study, we examined the effects of a number of putative MDCs on key aspects of pancreatic α-cell biology, including cell viability, gene expression profile, and glucagon secretion. Furthermore, by using the pancreatic α-cell line αTC1-9, we explore for the first time the biological applicability of this cell-based model as a potential tool for MDCs screening. In summary, we believe this will help to provide new insights into the field of endocrine disruption and unravel the effects and modes of action of MDCs at the pancreatic α-cell level.

## 2. Results

### 2.1. Bisphenols exert detrimental effects on pancreatic α-cell physiology

Bisphenol-A (BPA), bisphenol-S (BPS), and bisphenol-F (BPF) effects on pancreatic α-cell viability were assayed following 24, 48, or 72 h of exposure by using a combination of three classical and commonly used cell-cytotoxicity assays: the resazurin (RZ) assay, which evaluates mitochondrial metabolic activity; the neutral red uptake (NRU) assay, as an indicator of lysosomal activity; and the 5-carboxyfluorescein diacetate acetoxymethyl ester (CFDA-AM) assay, to determine plasma membrane integrity.

As shown in Figure 1A, the RZ assay indicated a slight decrease in pancreatic α-cell viability in BPA-treated cells (24 h) at the concentration range of 10 nM to 10 μM (reduction to 93–94% compared to control 100%). At the same doses, membrane integrity was also moderately reduced (reduction to 94–95% compared to control 100%). However, the NRU assay did not indicate cytotoxicity, except for a moderate effect at the 10 nM BPA concentration (93.90 ± 1.42%) (Figure 1A). In contrast, cell viability (measured as RZ test) in response to BPS treatment was slightly but significantly reduced only at the highest dose tested (10 μM) (Figure 1B), while membrane integrity was found to be decreased at all doses assayed except for 1 nM concentration (Figure 1B). As is the case for BPA, the NRU assay did not show any differences in cell viability between control and BPS-treated cells after 24 h (Figure 1B). Compared to BPA and BPS, BPF showed a more pronounced cytotoxicity effect. As shown in Figure 1C, decreased cell viability (RZ assay) was quantified at almost all BPF concentrations tested, from 1 nM to 10 μM. The maximum inhibitory effect was observed at 10 μM concentration (83.64 ± 1.19%) compared to the control (100.00 ± 1.13%). Membrane integrity was also found to be affected, with a minimum reduction effect observed at 1 nM (92.92 ± 1.55%) and a maximum effect at the highest dose of 10 μM (88.74 ± 1.27%) compared to the control (100.00 ± 0.98%) (Figure 1C). No effect was found in the NRU assay (Figure 1C). Overall, these data showed that BPA, BPS, and particularly BPF slightly impaired mitochondrial activity and membrane integrity, while lysosomal activity was unaffected.

**Figure 1.**
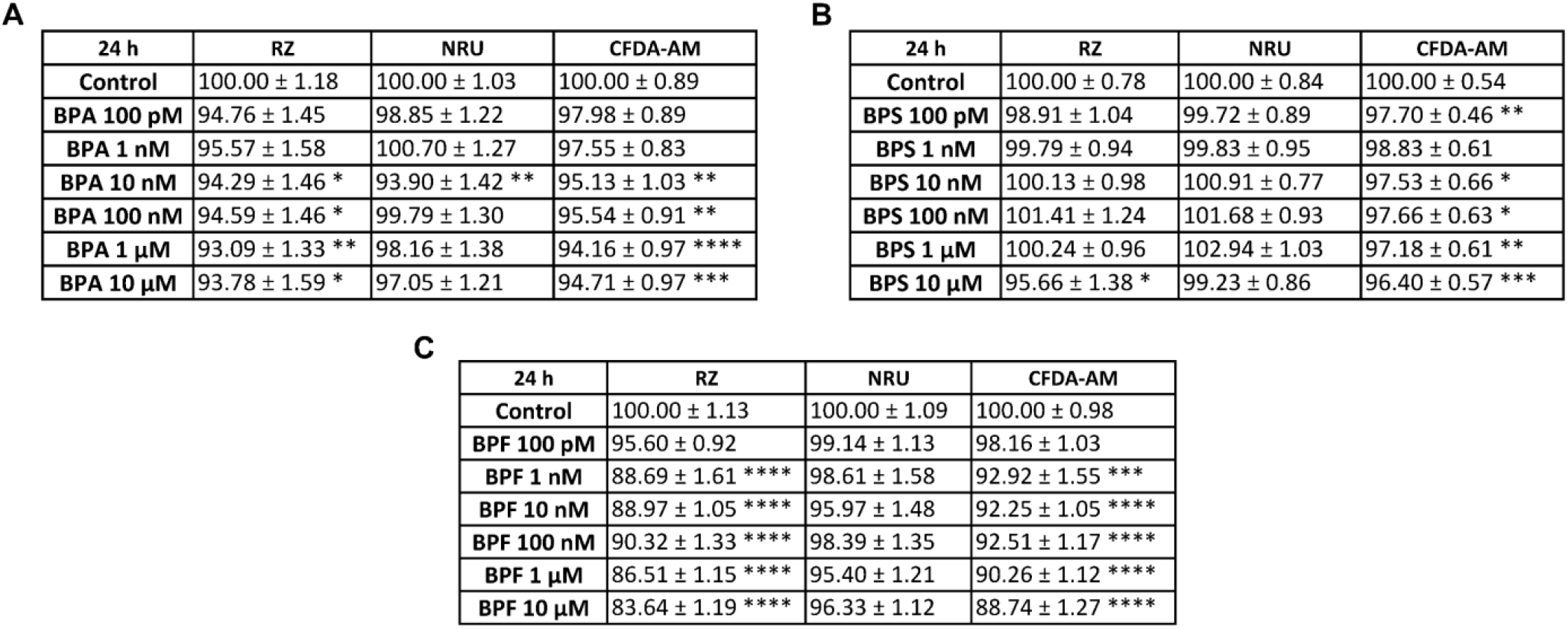
Viability of pancreatic αTC1-9 cells treated for 24 h with different concentrations of bisphenols. Cells were treated with a range of concentrations (100 pM–10 μM) of (A) BPA, (B) BPS or (C) BPF. Viability was evaluated with RZ, NRU and CFDA-AM assays for each compound. Results are expressed as % of the solvent control (Control=100%). Data are represented as mean ± SEM of (A) n= 4, (B) n=5, and (C) n=4 independent experiments. * vs. Control; * *p* < 0.05, ** *p* < 0.01, *** *p* < 0.001, **** *p* < 0.0001 by one-way ANOVA followed by Dunnett’s post hoc test or Kruskal-Wallis followed by Dunn’s post hoc test.

Cell viability was also explored after 48 and 72 h treatment with different BPA, BPS, or BPF concentrations (100 pM–10 μM). The most remarkable changes for BPA were found at 48 h, while no cytotoxic effects were observed at 72 h (Supplemental Table S1 and Table S2). At 48 h, mitochondrial (1, 100 nM and 1 μM) and lysosomal (1 and 10 μM) activity, as well as membrane integrity (100 nM, 1 and 10 μM), were slightly reduced in BPA-treated α-cells compared to the control. On the contrary, 48 h treatment with BPS did not impact cell viability (Supplemental Table S1). At 72 h, a modest decrease in cell viability was measured using RZ (1, 10 nM and 10 μM) and NRU assays (1 nM). No effects were found at the level of membrane integrity (Supplemental Table S2). As observed in the 24 h treatment, BPF provoked a more pronounced effect on cell viability. Following 48 h treatment, decreased mitochondrial activity (1 nM–10 μM) was found, with a maximum effect at 10 μM dose (86.61 ± 1.27) (Supplemental Table S1). Similar effects were observed at 72 h, with a significant reduction at all doses tested (Supplemental Table S2). At both time points, 48 and 72 h, membrane integrity was also moderately reduced in response to BPF treatment at the concentration range of 1 nM to 10 μM (Supplemental Table S1, S2). No effects were observed in the NRU assay, except for a slight reduction at 48 h (10 μM) (Supplemental Table S1).

We next analysed whether the treatment with bisphenols could alter the expression of genes relevant to pancreatic α-cell function and identity. In particular, the expression pattern of genes encoding for glucagon (*Gcg*); the enzyme glucokinase (*Gck*), which is critical to glucose sensing, the glucose transporter for α-cells *Glut1*; as well as different transcription factors that are key for the maintenance of α-cell identity and function, such as aristaless-related homeobox (*Arx)*, v-maf avian musculoaponeurotic fibrosarcoma oncogene homolog B (*MafB)*, and forkhead box O1 (*Foxo1)*, were explored. The main change observed in response to BPA treatment was a reduction in *Gcg* gene expression, which was significant at 100 nM and 1 μM concentrations (Figure 2A). On the contrary, both *Gck* and *MafB* gene expression was found to be increased at 100 nM dose, while at 1 μM, *MafB* was decreased (Figure 2B, E). Furthermore, *Foxo1* expression was elevated at the highest BPA dose tested (Figure 2F). No significant effects on *Glut1* or *Arx* genes (Figure 2C, D) were found. In a similar manner, BPS exposure resulted in decreased *Gcg* expression compared to the control (Figure 3A), although, unlike BPA, the effect was already visible at the lowest dose tested (100 pM) and was maintained over a wider dose range, including 10, 100 nM and 10 μM concentrations. Decreased expression of *Gck* (1 and 10 μM) and *Foxo1* (100 pM and 10 nM) was also observed in BPS-treated α-cells (Figure 3B, F). No changes were observed in *Glut1* (Figure 3C), *Arx* (Figure 3D), or *MafB* (Figure 3E) gene expression levels in response to BPS treatment. BPF did not promote any significant effect on *Gcg* (Figure 4A), *Gck* (Figure 4B) or *Glut1* expression (Figure 4C), although *Arx* (Figure 4D) and *MafB* (Figure 4E) gene expression was attenuated at 1 nM and 10 μM, and 100 pM and 1 μM doses, respectively. In addition, BPF-treated cells showed a decreased expression of *Foxo1* at 100 nM concentration (Figure 4F).

**Figure 2.**
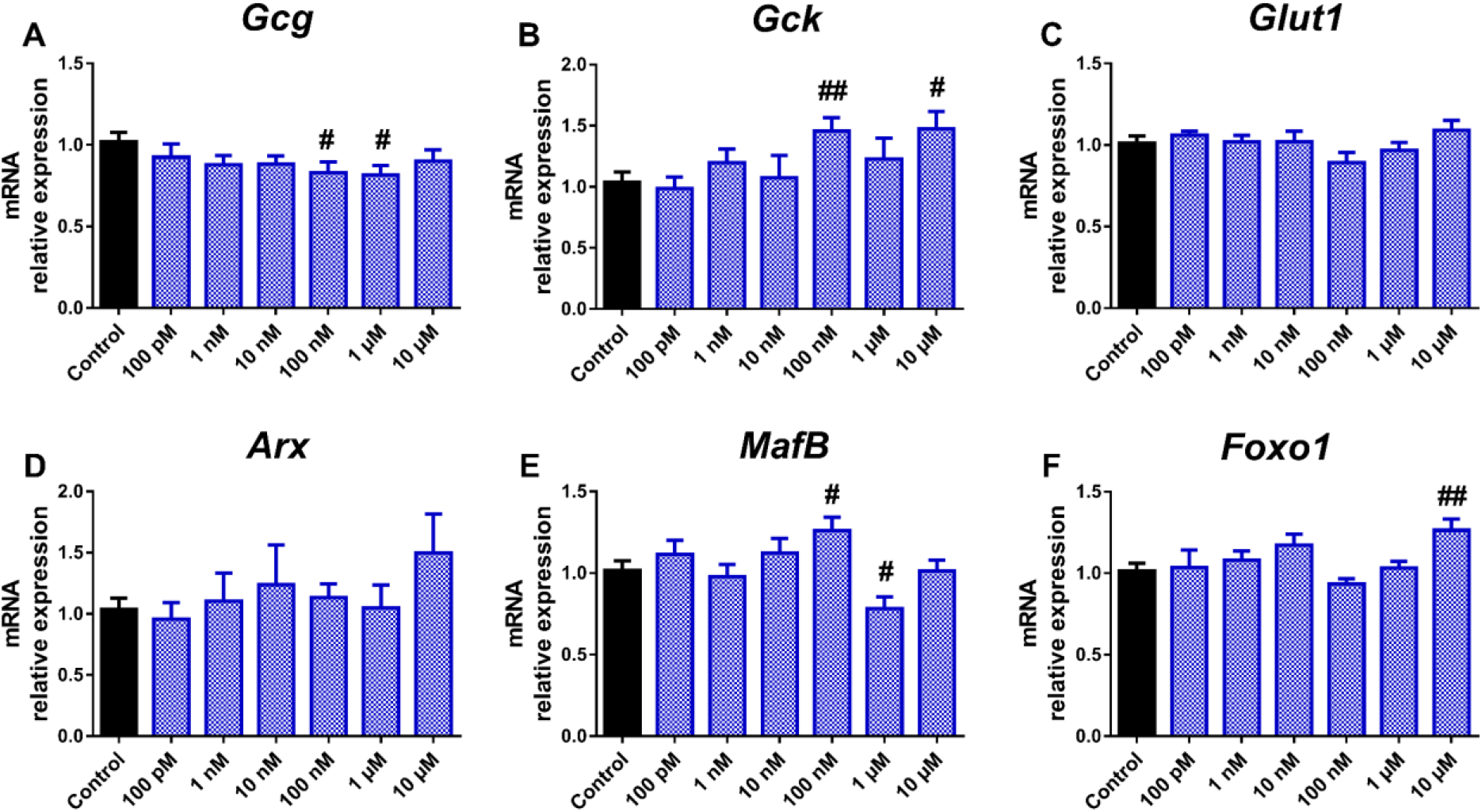
BPA effects on the expression of genes involved in pancreatic α-cell function and identity. mRNA expression of (A) *Gcg*, (B) *Gck*, (C) *Glut1*, (D) *Arx*, (E) *MafB*, and (F) *Foxo1* was measured in αTC1-9 cells treated for 24 h with different BPA concentrations (100 pM–10 μM). Data are represented as mean ± SEM of n= 3–4 independent experiments. # vs. Control; # *p* < 0.05, ## *p* < 0.01 by the Student’s t test.

**Figure 3.**
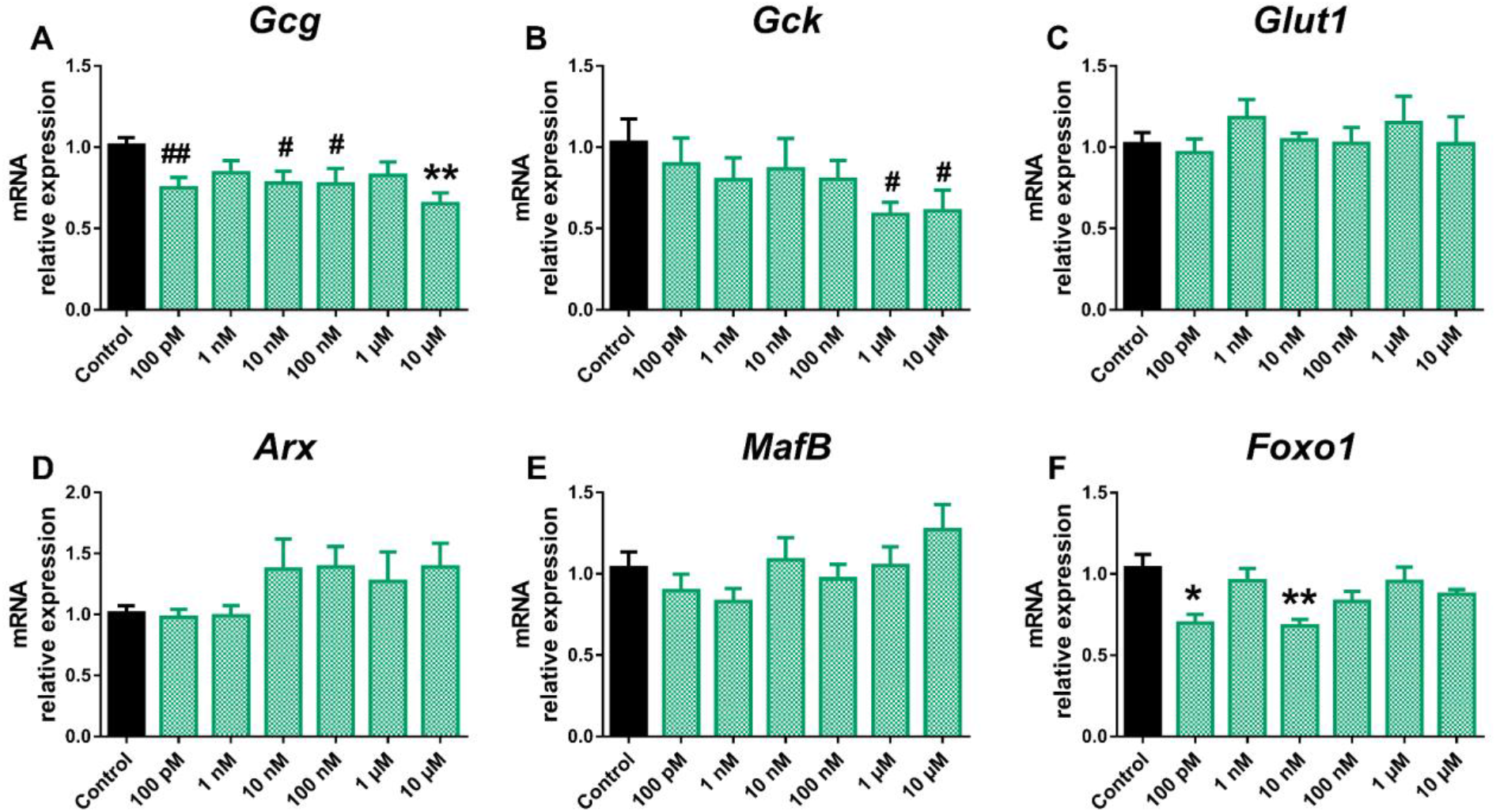
BPS effects on the expression of genes involved in pancreatic α-cell function and identity. mRNA expression of (A) *Gcg*, (B) *Gck*, (C) *Glut1*, (D) *Arx*, (E) *MafB*, and (F) *Foxo1* was measured in αTC1-9 cells treated for 24 h with different BPS concentrations (100 pM–10 μM). Data are represented as mean ± SEM of n= 3–4 independent experiments. * vs. Control; * *p* < 0.05, ** *p* < 0.01 by one-way ANOVA followed by Dunnett’s post hoc test or Kruskal-Wallis followed by Dunn’s post hoc test. # vs. Control; # *p* < 0.05, ## *p* < 0.01 by the Student’s t test.

**Figure 4.**
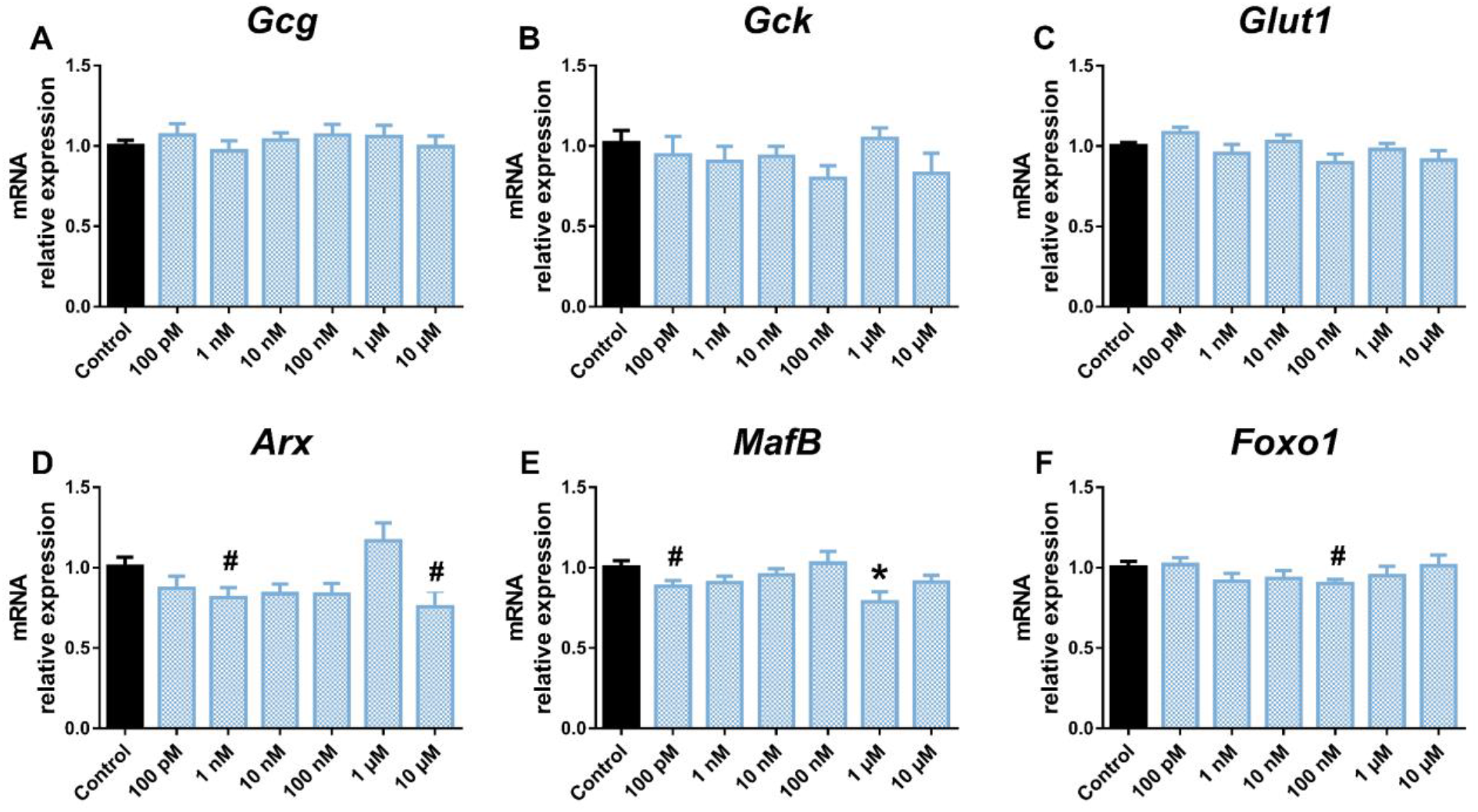
BPF effects on the expression of genes involved in pancreatic α-cell function and identity. mRNA expression of (A) *Gcg*, (B) *Gck*, (C) *Glut1*, (D) *Arx*, (E) *MafB*, and (F) *Foxo1* was measured in αTC1-9 cells treated for 24 h with different BPF concentrations (100 pM–10 μM). Data are represented as mean ± SEM of n= 3 independent experiments. * vs. Control; * *p* < 0.05 by one-way ANOVA followed by Dunnett’s post hoc test. # vs. Control; # *p* < 0.05 by the Student’s t test.

To decipher whether the above changes were related to altered pancreatic α-cell function, glucagon secretion was assayed in cells treated with vehicle, BPA, BPS, or BPF for 24 h. Glucagon secretion was analysed after incubating the cells with 0.5 mM glucose, 16 mM glucose, or 0.5 mM glucose plus 10 nM insulin. As expected in control conditions, cells exhibited a maximum glucagon release at a low glucose concentration (0.5 mM), as this is the major physiological stimulus for glucagon release, while significant decreases were observed at both high glucose (16 mM) and low glucose plus insulin, stimuli that act as negative regulators of α-cell function. In turn, all analysed bisphenols led to a significant inhibitory effect on glucagon secretion. However, we found significant differences in the effective doses. The BPA inhibitory response was only significant at 10 nM concentration (Figure 5A); however, BPS promoted a marked decline in glucagon release at all doses assayed with a maximum effect at the lowest one, 1 nM (Figure 5B). BPF effects were relatively less pronounced compared to BPA and BPS. Of note, BPF displayed a non-monotonic dose–response behaviour with a reduction at 1 and 100 nM concentrations but no effect at 10 nM (Figure 5C).

**Figure 5.**
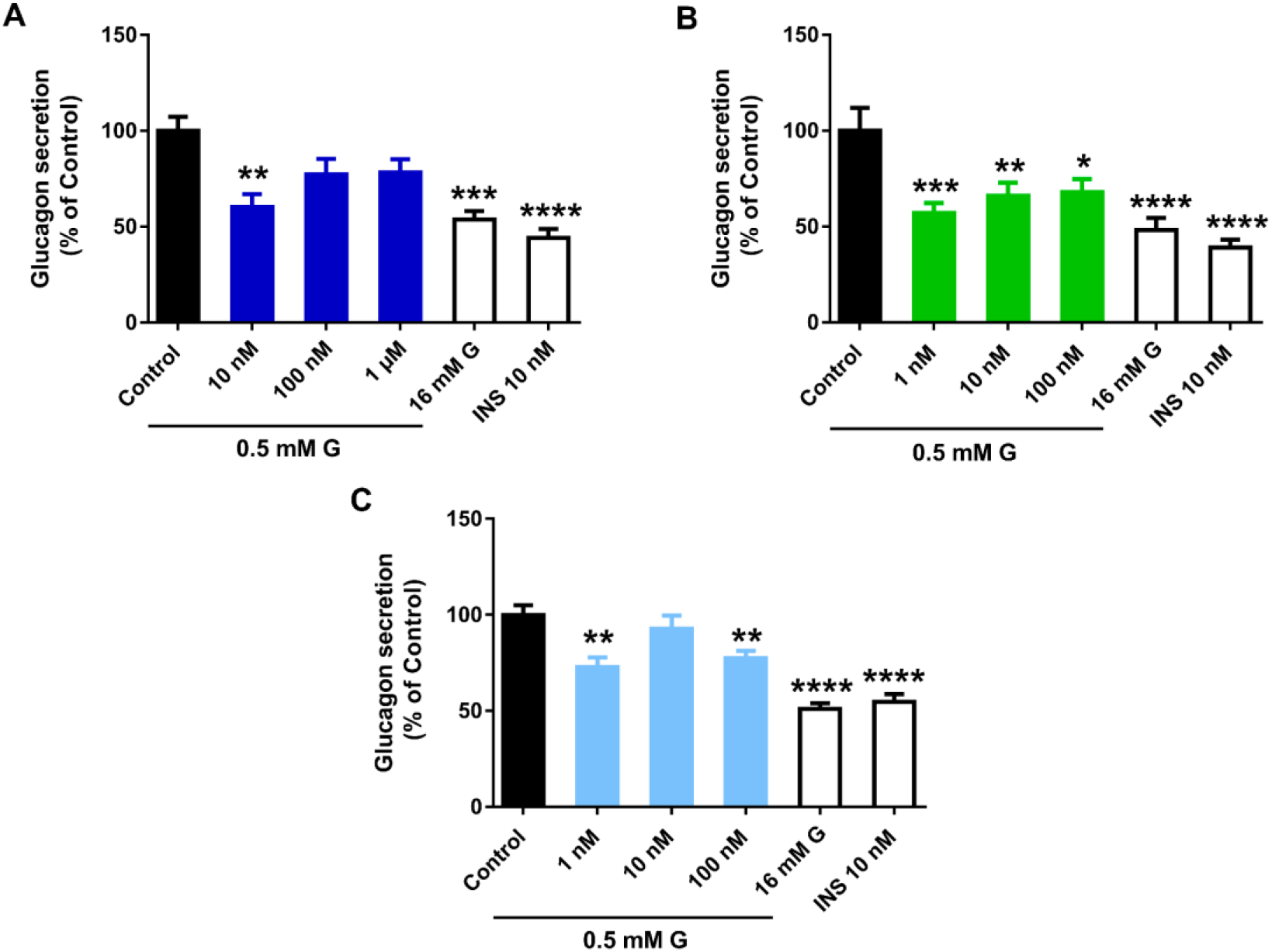
Bisphenols treatment disrupts pancreatic α-cell function. Effects of (A) BPA, (B) BPS and (C) BPF on glucagon secretion in αTC1-9 cells treated for 24 h. Glucagon release in control and bisphenol-treated cells was measured in response to 0.5 mM glucose (0.5 mM G). Glucagon secretion in response to 16 mM glucose (16 mM G) and 0.5 mM G plus 10 nM insulin (INS 10 nM) conditions were also included as additional controls. Data are represented as mean ± SEM of (A) n= 6, (B) n= 6, and (C) n=3 independent experiments. * vs. Control; * *p* < 0.05, ** *p* < 0.01, *** *p* < 0.001, **** *p* < 0.0001 by one-way ANOVA followed by Dunnett’s post hoc test or Kruskal-Wallis followed by Dunn’s post hoc test.

### 2.2. Di(2-ethylhexyl) phthalate (DEHP) impaired pancreatic α-cell function with no major changes in cell viability or gene expression

DEHP effects on pancreatic α-cell viability were assayed after 24, 48, or 72 h of exposure. As illustrated in Figure 6A, no significant cytotoxic effects of DEHP were observed in pancreatic α-cells treated for 24 h in any of the three cell cytotoxicity tests assayed, except for a slight decrease in mitochondrial activity at 100 nM (95.64 ± 1.32%) and membrane integrity at 1 μM (95.61 ± 0.90%) concentrations. At longer times of exposure, the decline in mitochondrial activity and membrane integrity was slightly more pronounced (Supplemental Table S1, S2). At 48 h, the RZ assay indicated a similar reduction in cell viability in the dosage range of 1 nM–1 μM with a maximum effect at 1 μM dose (93.25 ± 1.42%) (Supplemental Table S1). At 72 h, the reduction was up to 91.38 ± 1.07% compared to the control (100.00 ± 1.39%) in response to 10 nM DEHP (Supplemental Table 2). Membrane integrity was also altered with a modest decrease at 48 h (10 nM–10 μM) and 72 h (1 nM–10 μM). In both cases, a maximal reduction effect was found at 1 μM concentration (Supplemental Table S1, S2). No effects were observed in the lysosomal activity at any time points assayed (Figure 6A and Supplemental Table S1, S2). At the gene-expression level, no significant effects were observed in DEHP-treated cells except for increased *Gcg* gene expression at 100 pM, 100 nM, and 10 μM, and a tendency toward increased expression at the rest of the doses tested (Figure 6B). In addition, a reduction in *Glut1* was quantified, although the effect was only statistically significant at the highest concentration assayed, 10 μM (Figure 6B). Despite the enhanced expression found on the *Gcg* gene, glucagon secretion in response to low glucose was diminished when cells were treated with DEHP for 24 h. The effect was visible at 100 nM and 1 μM but not at a lowest dose, 10 nM (Figure 6C).

**Figure 6.**
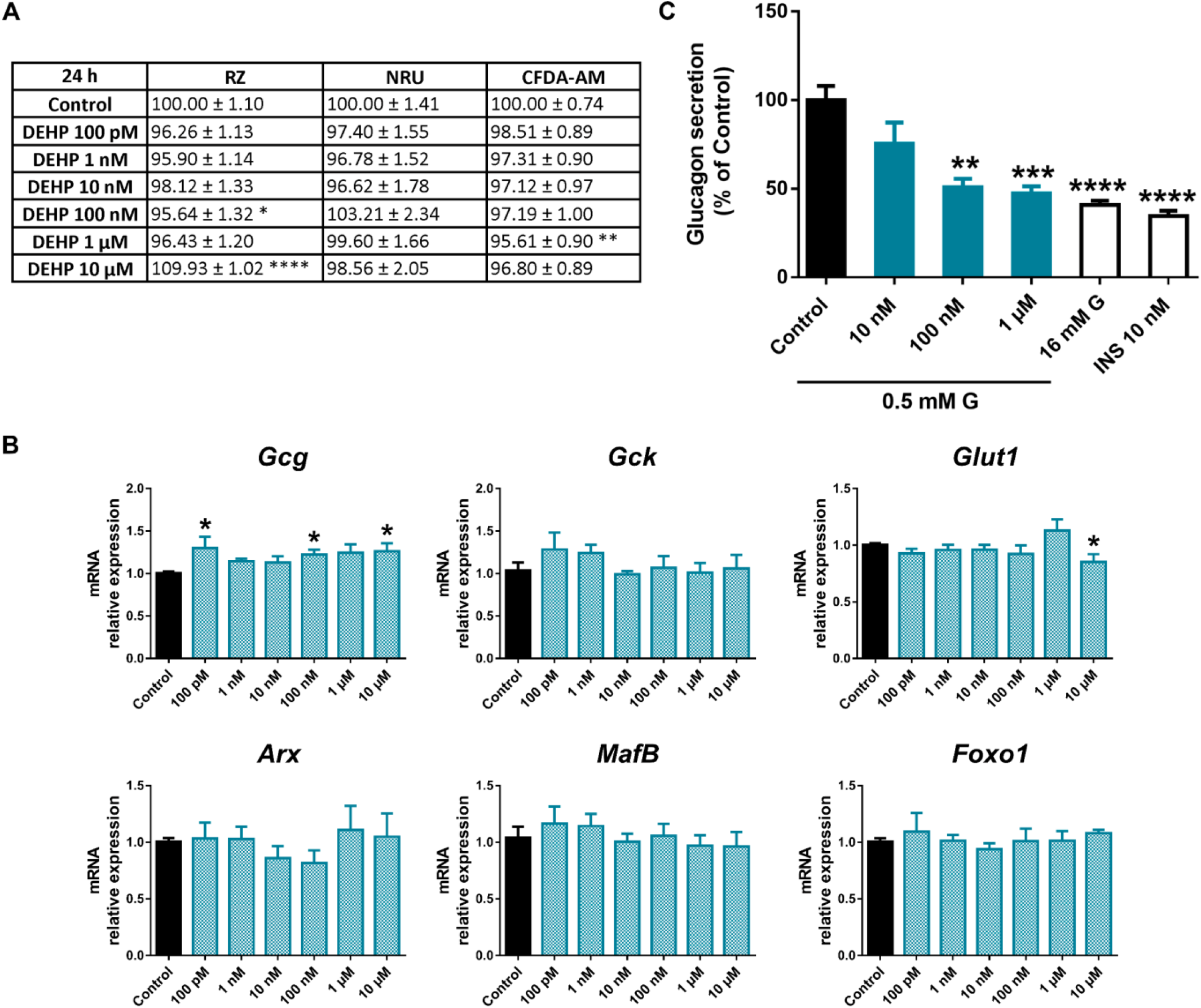
DEHP effects on pancreatic α-cells. A) Viability of pancreatic αTC1-9 cells treated for 24 h with different concentrations of DEHP (100 pM–10 μM) as evaluated by RZ, NRU and CFDA-AM assays. Results are expressed as % of the solvent controls (Control=100%). B) mRNA expression of *Gcg, Gck, Glut1, Arx, MafB*, and *Foxo1* in control and DEHP-treated αTC1-9 cells for 24 h. C) Glucagon secretion in αTC1-9 cells treated for 24 h with DEHP. Data are represented as mean ± SEM of (A) n= 3, (B) n= 3, (C) n= 5 independent experiments. * vs. Control; * *p* < 0.05, ** *p* < 0.01, *** *p* < 0.001, **** *p* < 0.0001 by one-way ANOVA followed by Dunnett’s post hoc test or Kruskal Wallis followed by Dunn’s post hoc test.

### 2.3. Perfluorooctanesulfonic acid (PFOS) did not compromise pancreatic α-cell function

PFOS treatment led to a modest decline in pancreatic α-cell viability, as RZ, CFDA-AM and NRU test assays indicated, although some differences in the effective dose and exposure time were observed. Mitochondrial metabolic activity was reduced in response to a wide range of PFOS concentrations after 24 (10 nM–1 μM) (Figure 7A) and 48 h (1 nM–10 μM) treatment (Supplemental Table S1), while at 72 h, the effect was only significant at 10 nM (Supplemental Table S2). Lysosomal activity was also affected at 10 nM PFOS, both at 24 (Figure 7A) and 48 h time points (Supplemental Table S1), and to a wider extent at 72 h (10 nM, 1 and 10 μM) (Supplemental Table S2). Membrane integrity was reduced at 10, 100 nM and 10 μM concentrations at 24 h (Figure 7A) and in the dose range of 10 nM to 10 μM, and 10 nM to 1 μM at 48 h and 72 h, respectively (Supplemental Table S1, S2). Upregulated expression of *Gck, Glut1, Arx*, and *Foxo1* was found at various PFOS concentrations (10 nM, 1 μM, 100 pM, and 10 μM, respectively) (Figure 7B). Despite the above-described changes, no significant impact on glucagon secretion was observed in response to PFOS treatment (Figure 7C).

**Figure 7.**
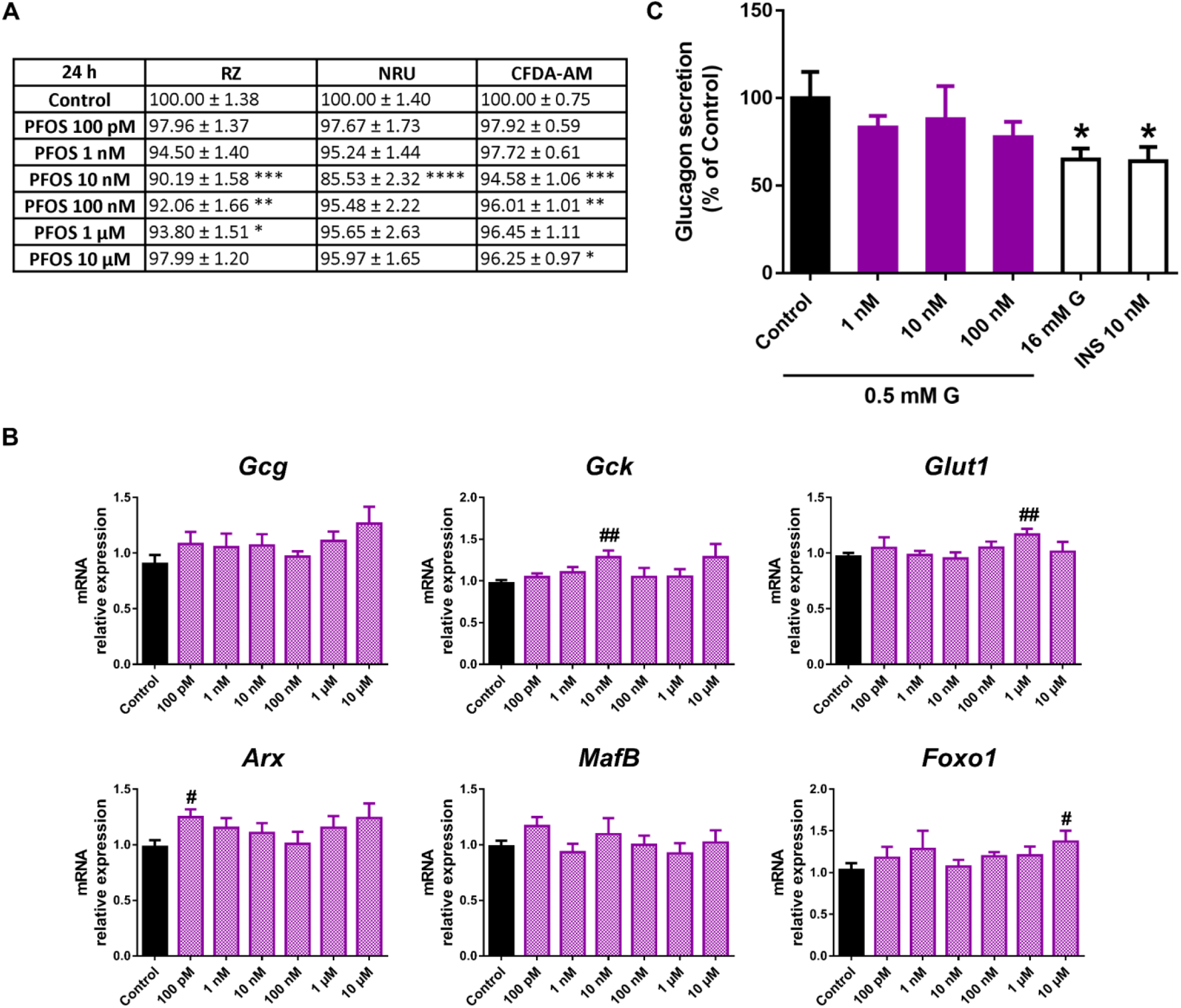
PFOS effects on pancreatic α-cells. A) Viability of pancreatic αTC1-9 cells treated for 24 h with different concentrations of PFOS (100 pM–10 μM) as evaluated by RZ, NRU and CFDA-AM assays. Results are expressed as % of the solvent controls (Control=100%). B) mRNA expression of *Gcg, Gck, Glut1, Arx, MafB*, and *Foxo1* in control and PFOS-treated αTC1-9 cells for 24 h. C) Glucagon secretion in αTC1-9 cells treated for 24 h with PFOS. Data are represented as mean ± SEM of (A) n= 3, (B) n= 3, and (C) n= 3 independent experiments. * vs. Control; * *p* < 0.05, ** *p* < 0.01, *** *p* < 0.001, **** *p* < 0.0001 by one-way ANOVA followed by Dunnett’s post hoc test or Kruskal Wallis followed by Dunn’s post hoc test. # vs. Control; # *p* < 0.05, ## *p* < 0.01 by the Student’s t test.

### 2.4. Cadmium chloride (CdCl_2_) effects on pancreatic α-cell function and viability

Following 24 h of CdCl_2_ exposure, no effects on cell viability were reported (Figure 8A). Similar results were found at 48 h and 72 h, despite a modest decrease in lysosomal activity at 48 h (100 nM, 1 and 10 μM) and mitochondrial activity at 72 h (10 μM) (Supplemental Table S1, S2). No significant changes in gene expression were quantified either, except for enhanced *Gcg* expression at 10 μM CdCl_2_ (Figure 8B). However, a marked decrease in glucagon secretion in response to low glucose concentration was found at all the concentrations assayed (1–100 nM) (Figure 8C).

**Figure 8.**
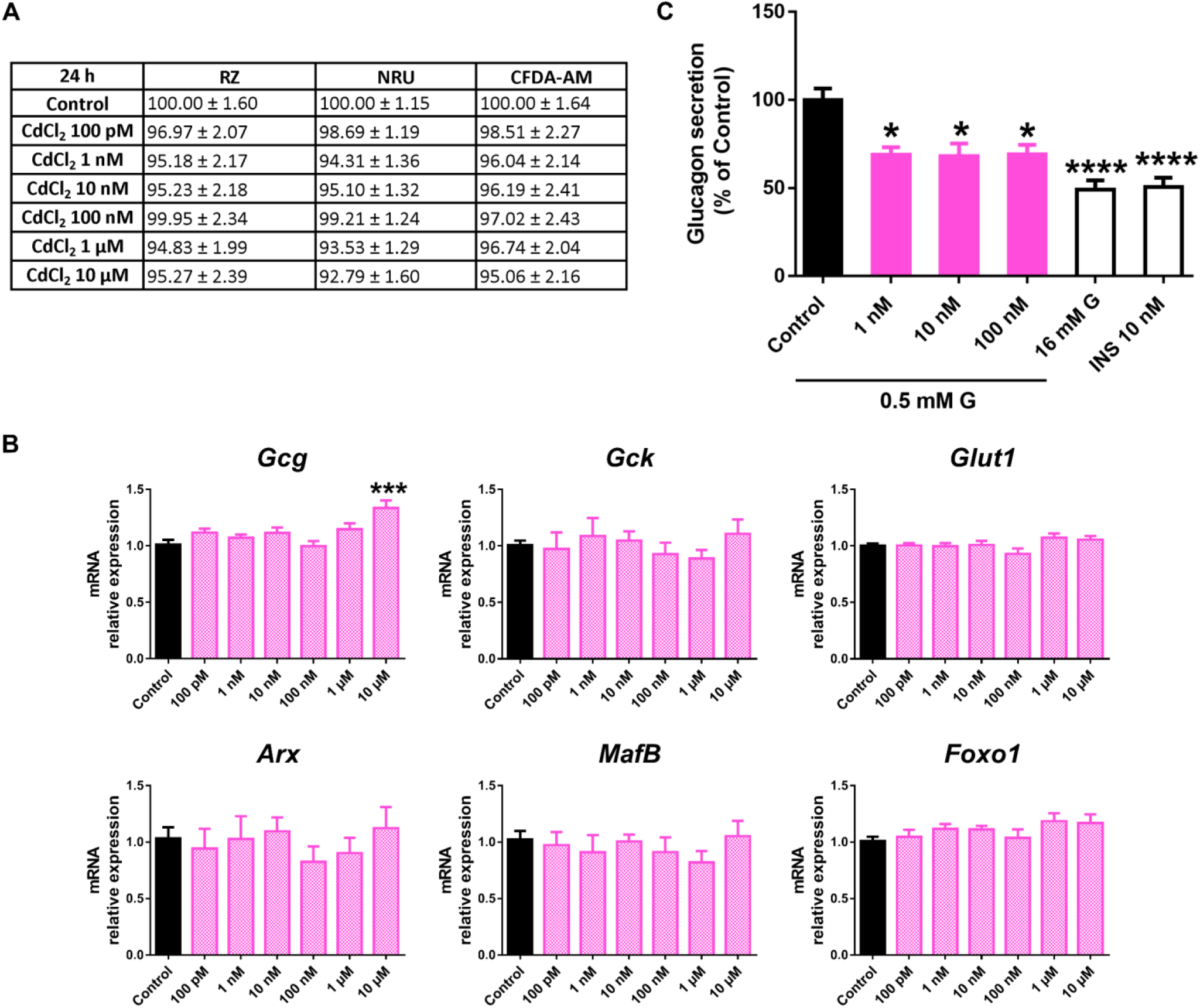
CdCl_2_ on pancreatic α-cells. A) Viability of pancreatic αTC1-9 cells treated for 24 h with different concentrations of CdCl_2_ (100 pM–10 μM) as evaluated by RZ, NRU and CFDA-AM assays. Results are expressed as % of the solvent controls (Control=100%). B) mRNA expression of *Gcg, Gck, Glut1, Arx, MafB*, and *Foxo1* in control and CdCl_2_-treated αTC1-9 cells for 24 h. C) Glucagon secretion in αTC1-9 cells treated for 24 h with CdCl_2_. Data are represented as mean ± SEM of (A) n= 3, (B) n= 3, and (C) n= 6 independent experiments. * vs. Control; * *p* < 0.05, *** *p* < 0.001, **** *p* < 0.0001 by one-way ANOVA followed by Dunnett’s post hoc test or Kruskal Wallis followed by Dunn’s post hoc test.

### 2.5. The impact of dichlorodiphenyldichloroethylene (DDE) on pancreatic α-cell physiology

The 24 h DDE treatment promoted a modest decrease in mitochondrial activity (Figure 9A). The effect was only significant at concentrations in the micromolar range; on the contrary, the effect was visible at all doses tested (100 pM–10 μM) at 48 h (Supplemental Table S1) and 100 pM, 100 nM, and 1, 10 μM at 72 h (Supplemental Table S2). Membrane integrity was also consistently reduced at most time points of exposure assayed, with a maximum effect at 10 μM dose following 24 (95.94 ± 0.63%) (Figure 9A), 48 (90.60 ± 1.19%) (Supplemental Table S1), and 72 (96.28 ± 0.59%) (Supplemental Table S2) h treatment. In turn, the NRU assay manifested only a moderate decrease in lysosomal activity at the lowest dose, 100 pM, at 24 h (Figure 9A), while no differences were observed at any other dose or time of treatment in DDE-cells compared to controls, except for a slight increase at 10 μM (48 h) (Supplemental Table S1, S2). At the gene-expression level, we found a dramatic reduction in *Glut1* mRNA levels, which was significant at all doses assayed from 100 pM to 10 μM (Figure 9B). In addition, some minor changes were quantified, including the enhanced expression of *Gck* and *Arx* at 100 pM and 1 μM doses, respectively (Figure 9B). Finally, a slight decrease in glucagon secretion was found, although the inhibitory effect was only significant at 1 nM concentration (Figure 9C).

**Figure 9.**
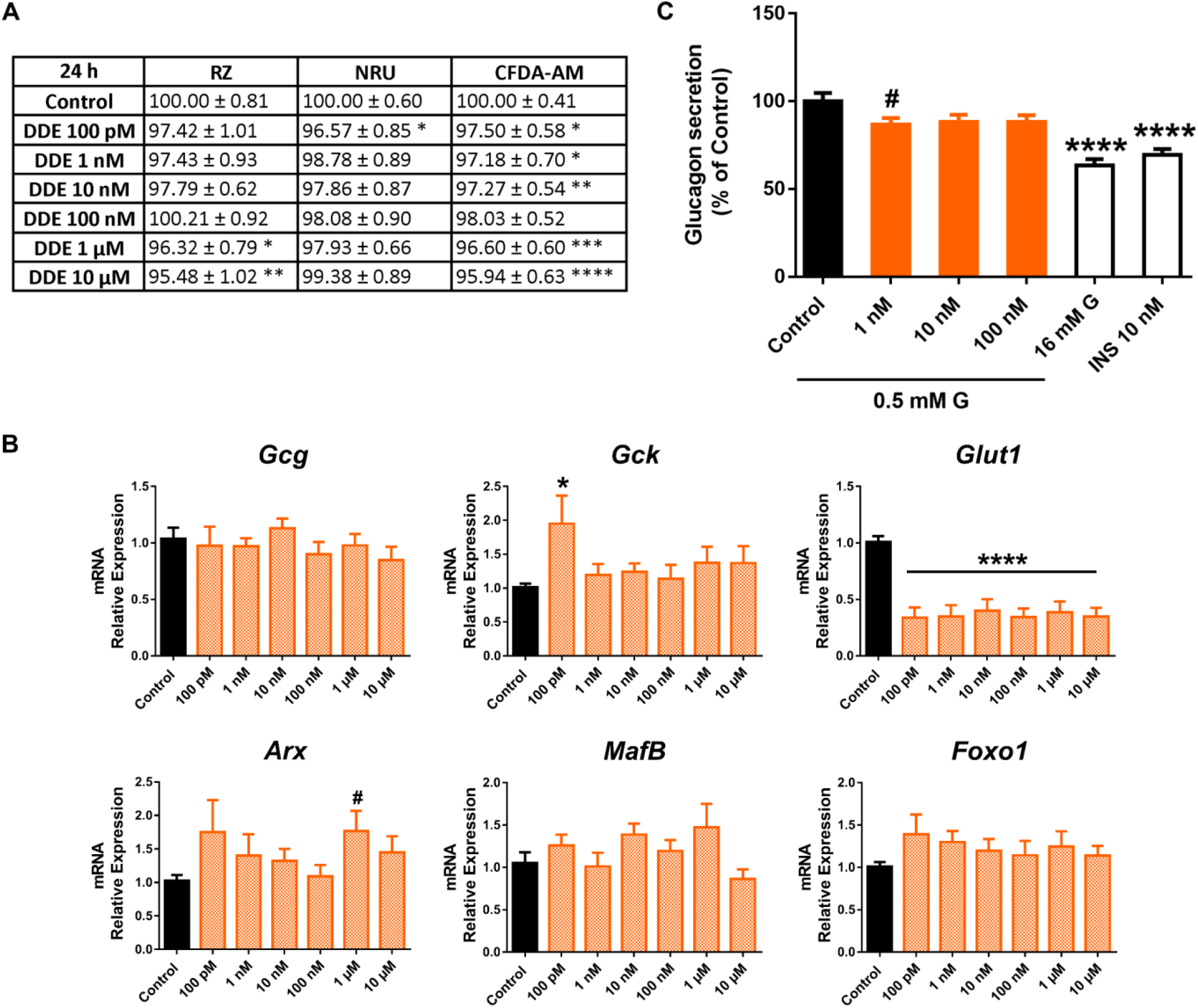
DDE effects on pancreatic α-cells. A) Viability of pancreatic αTC1-9 cells treated for 24 h with different concentrations of DDE (100 pM–10 μM) as evaluated by RZ, NRU and CFDA-AM assays. Results are expressed as % of the solvent controls (Control=100%). B) mRNA expression of *Gcg, Gck, Glut1, Arx, MafB*, and *Foxo1* in control and DDE-treated αTC1-9 cells for 24 h. C) Glucagon secretion in αTC1-9 cells treated for 24 h with DDE. Data are represented as mean ± SEM of (A) n= 5, (B) n= 3, (C) n= 6 independent experiments. * vs. Control; * *p* < 0.05, ** *p* < 0.01, *** *p* < 0.001, **** *p* < 0.0001 by one-way ANOVA followed by Dunnett’s post hoc test or Kruskal Wallis followed by Dunn’s post hoc test. # vs. Control; # *p* < 0.05 by the Student’s t test.

## 3. Discussion

In the present study, we have screened the effects of a number of paradigmatic EDCs on key aspects of pancreatic α-cell physiology, including cell viability, gene expression, and glucagon secretion, using an in vitro model, the mouse α-cell line αTC1-9. We selected seven MDCs representing five major toxic classes of chemicals used as plasticizers, pesticides, or heavy metals. The αTC1-9 cells are considered a robust and suitable cell model for studying pancreatic α-cell function as they replicate a phenotype consistent with that of differentiated α-cells in normal islets, and properly resemble their physiology [20-23]. As a proof of concept, αTC1-9 cells produce the α-cell-specific hormone glucagon but not insulin or any other islet hormones [20, 24], express transcription factors that are key for α-cell specification (*Nkx2*.*2, FoxA2, Arx*, and *Pax6*), but not for β-cell specification (*Pdx1, Pax4, Nkx6*.*1* and *HNF4α*, and *MafA*) [20], and present a large number of common transcriptional regulatory elements similar to that of primary mouse and human α-cells [25]. Besides the above-mentioned points, it is important to note that important challenges must be faced when working with primary pancreatic α-cells. The main reason for this is the scarcity of this cellular type in islets, particularly in rodents, together with technical limitations since the appropriate methodology is needed to purify and identify α-cells, including the use of single-cell technologies or reporter animal models [26-30]. Therefore, using an immortalized cell line such as αTC1-9 offers a relatively simple and readily reproducible approach to studying pancreatic α-cell function.

We found that among the MDCs tested, those with the greatest effect on α-cell viability were BPA, BPF, and PFOS, while the rest of the compounds evaluated did not exert major cytotoxic effects. Cell viability was moderately affected in α-cells after 24 h of exposure as measured with the RZ assay. For BPF, the decline was around 11-16%, 6-7% for BPA, and in the case of PFOS, the reduction was approximately 7-10% within the nano- and micromolar range in all cases. These results indicate that mitochondrial metabolism could be impaired to some extent by the mentioned pollutants, as RZ reduction has been demonstrated to be highly correlated to oxygen consumption [31-33]. Similar results were found using the CFDA-AM assay with a maximum reduction effect on viability for BPF (8-12%) and around 5% for BPA and PFOS. Although significant, in all cases, cytotoxicity was relatively low, which may be attributed to the fact that glucagon-producing α-cells showed enhanced protection against the toxic environment created by certain insults. The mechanism underlying this phenomenon is not fully understood, although some scientific evidence suggests that α-cells may face and adapt to long-term metabolic stress by expressing higher levels of the anti-apoptotic protein Bcl2l1 [34]. IL-6 has also been suggested to protect α-cells from apoptosis induced by metabolic stress and promote α-cell mass expansion during obesity as a compensatory response. However, prolonged elevated IL-6 circulating levels may lead to α-cell failure [35]. Additionally, the high expression of UCP2 in α-cells seems to play a cytoprotective role against different stressors [36]. The activation of these signalling pathways in α-cells upon exposure to MDCs needs further investigation.

Another important aspect to consider is whether or not MDCs may regulate pancreatic α-cell gene expression and if the functional outcomes described in this study may respond to the gene-expression pattern modulation. In this regard, pancreatic αTC1-9 cells were especially sensitive to bisphenols action. In particular, we found that BPS, at low doses, markedly impaired low-glucose-induced glucagon secretion. This was associated with the decreased expression of the glucagon gene in response to BPS at nanomolar doses (10 and 100 nM) but also at a lower (100 pM) and a higher dose (10 μM). In addition, *Foxo1* mRNA levels were reduced at 100 pM and 10 nM BPS concentrations. Similarly, BPF exposure promoted decreased glucagon secretion (1, 100 nM) and *Foxo1* gene expression (100 nM). These data are in accordance with previous findings indicating that *Foxo1* may behave as a positive regulation factor, as *Foxo1* silencing reduced glucagon gene expression in αTC1-9 cells [37]. Of note, orexin-A (OXA), a neuropeptide involved in the regulation of food intake and energy homeostasis, has been shown to decrease glucagon expression and secretion in a mechanism dependent on Foxo1 phosphorylation [38], a process which leads Foxo1 to be excluded from the nucleus and therefore inactivated [39]. Thus, we speculate that there may be a connection between the decrease in *Foxo1* expression and the decline in glucagon secretion in response to BPS and BPF treatment at least at 10 nM and 100 nM doses, respectively. Remarkably, BPF also promoted a decline in the expression of the *Arx* gene at 100 nM dose. This is of relevance as Arx is an important transcription factor that is critical in the differentiation of α-cells. In addition, it has been demonstrated that the ablation of *Arx* results in a loss of α-cells and a concomitant increase in β and δ cells [40-43], which may indicate that a lower expression of *Arx* might be connected with a decrease in the number and/or function of α-cells. Whether this is the case in BPF-treated αTC1-9 cells, merits further investigation.

BPA, like BPS and BPF, also promoted a decline in glucagon secretion, which was significant at 10 nM concentration, and manifested a consistent tendency toward diminished secretion at 100 nM and 1 μM concentrations. This effect was accompanied by downregulated mRNA levels of the glucagon gene at 100 nM and 1 μM BPA doses. In line with this, previous findings from our lab revealed that low doses of BPA can suppress low glucose-induced intracellular Ca^2+^ oscillations in α-cells, a signal which is key for triggering glucagon secretion [44].

Unlike what has been previously described for BPS and BPF, no major effects on *Foxo1* or *Arx* transcript levels were found in response to BPA treatment. However, an upregulation of *MafB* gene expression in BPA-treated cells was found. *MafB* is a transcription factor expressed in both pancreatic α and β-cells in the embryonic pancreas but becomes exclusively restricted to α-cells in the adult pancreas. Besides its importance in α-cell development, *MafB* has also been reported to promote glucagon gene expression and be important for the functional maintenance of adult α-cells [45, 46]. Here, we found that BPA (100 nM and 1 μM) treatment led to enhanced *MafB* expression, which might be a compensatory response to the downregulation of mRNA glucagon levels found at the same BPA concentrations.

Interestingly, when αTC1-9 cells were exposed to either DEHP or CdCl_2_, a marked decline in pancreatic α-cell function was also found, although higher doses were needed for DEHP to impair glucagon secretion. In contrast with bisphenols, the reduction in glucagon release was not accompanied by changes in the gene expression profile except for a slight increase in glucagon mRNA levels in the DEHP-treated cells.

In the case of DDE, the most significant effect was a dramatic reduction in mRNA levels of the *Glut1* gene, which was significant at all doses tested. Contrary to what might be expected, this was not accompanied by important changes in glucagon secretion except for a slight decrease at 1 nM DDE. This could be because, although Glut 1 has been traditionally considered the main glucose transporter in α-cells [47], the sodium– glucose transporters 1 and 2 (SGLT1 and SGLT2) have also been shown to be present in α-cells [48]. Recent findings have found that canagliflozin, an inhibitor of SGLT1, repressed glucagon secretion in αTC1-9 cells in a mechanism dependent on glucose transport and intracellular Ca^2+^ increase. In addition, a correlation between SGLT1 mRNA levels and glucagon release was reported. On the contrary, the inhibition of *Glut1* by phloretin resulted in enhanced glucagon secretion in mouse islets, indicating that Glut 1 and SGLT1 may regulate glucagon release oppositely [49]. It remains unclear whether or not DDE may also modulate SGLT1 expression and if the final effect on glucagon release depends on the joint action of both transporters. In addition, it should be noted that modulation of the intracellular sodium level also seems to be important for glucagon secretion [50, 51] and, therefore, not only glucose uptake regulation must be taken into account.

At the functional level, we found that bisphenols, DEHP, and CdCl_2_, at environmentally relevant doses, can promote an impairment in the glucose-induced glucagon-secretion process. To the best of our knowledge, this is the first study to evaluate the direct effects of MDCs on α-cell secretion. Using an in vitro cellular model, we performed a screen to search for individual chemical effects on α-cell physiology. The model, the αTC1-9 cell line, is not exempt from certain limitations, as it is a 2D rodent cellular model, and therefore it cannot accurately represent the cellular proximity and anatomical structure arrangement within the islets of Langerhans. Nevertheless, it should be emphasized that 2D in vitro models constitute a particularly suitable approach for systematic and reproducible studies on specific cell types, especially for chemical testing. In addition, they represent an animal-free tool that is more cost-effective and easier to handle compared to 3D cell culture or animal models. As such, we believe that the current system offers a rapid assay platform for MDCs as a first-line screening of pancreatic α-cell biology.

Despite its obvious importance, there is a compelling need for further chemical testing as a second step under in vivo assays. To date, in vivo studies interrogating the impact of MDCs on α-cell physiology are still scarce, and in most cases, they draw contradictory conclusions. The discrepancy between studies may be attributed to different timings of exposure, dosing, and species-specificity, among other factors. For example, some investigations have shown that male zebrafish exposed to BPS for 28 d exhibited decreased glucagon gene expression [52], as was the case in flatfish treated with PFOS for 96 h [53]. In line with this, MEHP, a bioactive metabolite of DEHP, promoted decreased α-cell area in zebrafish embryos [54]. On the contrary, developmental exposure to BPA in mice resulted in increased glucagon expression and number of α-cells [55], while other authors did not find any effect on α-cell mass in the BPA-treated offspring [56]. Decreased α-cell proportion was observed in adult BPA-treated rats [57]. Glucagon levels were not found to be significantly changed either in response to DEHP [58, 59], PFOS [60], or CdCl_2_ [61] treatment. Nevertheless, as previously mentioned, studies are limited, and the in vivo investigation needs to be further extended.

Previous work from our group has demonstrated that most of the assayed chemicals also display direct and detrimental effects in both murine and human pancreatic β-cells [62]. In particular, we found that BPA, BPS, DEHP, PFOS, and CdCl_2_ were able to impair glucose-stimulated insulin secretion, insulin content, gene expression profile, and/or electrical activity in β-cells and that the effects were in general agreement with those described under in vivo conditions [62]. This is important if we consider that glucose homeostasis critically depends on the coordinated action of both pancreatic β and α-cells. The fact that these chemical compounds can alter the biology of both cellular systems highlights the significant harmful impact they can exert on glucose metabolism and, therefore, in the aetiology of metabolic disorders.

## 4. Conclusions

In recent decades, considerable research has been conducted to identify and understand the mechanisms underlying the adverse actions of EDCs. However, the scope of the findings made has been mainly limited to environmental chemicals with oestrogenic, androgenic, or steroidogenic activity, and comparatively, fewer studies have addressed other endocrine pathways. Of special interest are endocrine modalities with metabolic competence, as metabolism disorders have emerged as an important global health problem. Considering this, the impairment of metabolic pathways affecting the functional capacity of the endocrine pancreas is key. To date, a number of studies have explored the impact of MDCs on pancreatic β-cell function; however, little is known about the ability of these compounds to disrupt pancreatic α-cell biology, despite this cellular type playing a critical role in the maintenance of glucose homeostasis. The present study tries to shed some light on this aspect. The work presented highlights that most of the selected MDCs, at doses relevant to human exposure, have a moderate impact on α-cell viability but deleterious effects on glucagon secretion and gene-expression profile patterns. Our study also establishes the pancreatic α-cell line αTC1-9 as a potential tool for in vitro screening to identify MDCs that can directly affect pancreatic α-cells, and to elucidate the potential molecular mechanisms of these environmental pollutants. This latest aspect is particularly relevant as, to date, the pancreatic α-cell system is not covered by the current bioassay tests implemented by the European Union for regulatory purposes [19, 63].

## 5. Material and methods

### 5.1 Chemicals

BPA (Cat. No. 239658), BPS (Cat. No. 103039), BPF (Cat. No. B47006), DEHP (Cat. No. 36735), PFOS (Cat. No. 77283), CdCl_2_ (Cat. No. 202908), and DDE (Cat. No. 35487) were purchased from Sigma-Aldrich (Saint Louis, MO, USA). All the chemicals were dissolved in dimethyl sulfoxide (DMSO) to prepare the stock solution, except for CdCl_2_, which was dissolved in water.

### 5.2 Cell culture

αTC1-9 cells (American Type Cultures Collection (ATCC) CRL-2350; Barcelona, Spain) were cultured and passaged as previously described [22]. Briefly, cells were cultured in DMEM without phenol red (Sigma-Aldrich, Saint Louis, MO, USA) containing 16 mM glucose, 19 mM NaHCO_3_, 15 mM HEPES, 2 mM L-glutamine (Gibco, Paisley, UK), 0.1 mM non-essential amino acids (Gibco, NY, USA), 100 units/mL penicillin and 100 μg/mL streptomycin (Thermo Fisher Scientific, Waltham, MA, USA), and 10% fetal bovine serum (FBS) (HyClone, GE Healthcare Life Sciences, Logan, UT, USA). For EDC treatment, FBS was replaced by charcoal dextran-treated FBS (HyClone, GE Healthcare Life Sciences, Logan, Utah, USA). Cells were incubated at 37 °C with 5% CO_2_ and discarded after passage 25.

### 5.3 Cell viability

αTC1-9 cells were seeded at a density of 25 × 10^3^ cells per well in black 96-well plates (Corning Incorporated, Kennebunk, ME, USA) 72-96 h before the treatment. Then, the medium was replaced, and cells were treated with the different EDCs for 24, 48, or 72 h, as indicated in the figure legends. Medium was replaced every 24 h. Cell viability was measured by using a combination of three different indicator dyes: RZ (Thermo Fisher Scientific, Waltham, MA, USA), NR (Sigma-Aldrich, Saint Louis, MO, USA), and CFDA-AM (Thermo Fisher Scientific, Waltham, MA, USA). After the incubation period, cells were washed with phosphate-buffered saline (PBS). Then, cells were treated with a solution containing RZ (5% v/v) and CFDA-AM (4 μM) prepared in serum-free DMEM for 40 min. Fluorescence was measured using a fluorescence plate reader (POLARstar Omega, BMG Labtech). RZ is converted by dehydrogenase activity into resorufin (excitation 530-570 nm, emission 590-620 nm); CFDA-AM is cleaved by esterases and retained in the cells with intact membranes as a fluorescent product (excitation 485 nm, emission 520 nm). After the incubation period, solution containing RZ and CFDA-AM dye was removed, and cells were rinsed with PBS. Then, cells were incubated with NR solution for 2 h (0.005% w/v). NR dye solution was aspirated from the wells, and NR destaining solution (1% acetic acid–50% ethanol) was added after the cells were rinsed twice with PBS. NRU was quantified by measuring absorbance at 540 and 690 nm (background) using a microplate reader (Biotek EON). 10% DMSO was used as positive control for cellular damage. The results are expressed as percentages (%) of the readings in the control wells.

### 5.4 Gene expression

Cells were seeded in 24-well plates at a density of 150-200 × 10^3^ cells/well. RNA was extracted using a commercial kit (Extractme RNA & DNA Kit; Blirt, Poland) according to the manufacturer’s instructions. After completing the extraction process, the RNA was reverse-transcribed using the High Capacity cDNA Reverse Transcription kit (Applied Biosystems, Foster City, CA, USA). Quantitative PCR assays were performed using the CFX96 Real Time System (Bio-Rad Laboratories, Hercules, CA, USA). Amplification reactions were carried out in medium containing a 200 nM concentration of each primer, 1 μL of cDNA, and IQ SYBR Green Supermix (Bio-Rad Laboratories, Hercules, CA, USA). Primers were designed between exons to avoid genomic cross-reaction. Samples were subjected to the following conditions: 30 s at 95 °C, 45 cycles (5 s at 95 °C, 5 s at 60 °C, and 10 s at 72 °C) and a melting curve of 65–95 °C. The resulting values were analysed with the CFX96 Real-Time System (Bio-Rad Laboratories, Hercules, CA, USA) and were expressed relative to the control values (2−ΔΔCT). Measurements were performed in duplicate and normalized against the geometric mean of the three housekeeping genes *Actb, Hprt*, and *Gapdh*. The primers used herein are listed in Supplemental Table S3.

### 5.5 Glucagon-secretion measurement

αTC1-9 cells were seeded in 24-well plates at a density of 125 × 10^3^ cells per well 72-96 h before the treatment. Cells were treated with the different EDCs for 24 h. After the treatment period, cells were preincubated with modified Krebs–Ringer medium containing 120 mM NaCl, 5.4 mM KCl, 1.2 mM KH_2_PO_4_, 1.2 mM MgSO_4_, 20 mM HEPES, 2.4 mM CaCl_2_, 5.6 mM glucose, and 0.1% BSA, pH 7.4 at 37 °C for 2 h. Then, cells were washed with Krebs (0 mM glucose) and incubated with Krebs–Ringer medium (0.5 mM glucose) for 30 min. Cells were washed again with Krebs without glucose and then incubated with 16 mM glucose for 30 min, or 0.5 mM glucose + 10 nM insulin. Supernatants were collected and used to measure glucagon secretion using ELISA (Mercodia, Uppsala, Sweden). To measure glucagon release, aprotinin (20 mg/L) (Sigma-Aldrich, Saint Louis, MO, USA) was included in all media. The total protein concentration was analysed using the Bradford dye method. Glucagon secretion was normalized by protein content and expressed as percentage of control 0.5 mM glucose.

### 5.6 Statistical Analysis

Statistical analysis was performed using GraphPad Prism 7.0 software (GraphPad Software, Inc., San Diego, CA, USA). Data are expressed as the mean ± SEM. To examine differences between groups, one-way ANOVA followed by Dunnett’s post hoc test or Student’s *t*-test was used when appropriate. When data did not pass the parametric test, Kruskal–Wallis followed by post hoc Dunn’s multiple comparison test was used. Statistical significance was set at *p* < 0.05 for all the analyses.

## Author Contributions

Conceptualization, R.A.-A., H.F. and P.A.-M.; formal analysis, R.A.-A., H.F., T.B.-B., S.S., I.Q. and P.A.-M.; investigation, R.A.-A., H.F., T.B.-B. and P.A.-M.; resources, P.A.-M., I.Q.; writing—original draft preparation, P.A.-M.; writing—review and editing, R.A.-A., H.F., T.B.-B., S.S., I.Q. and P.A.-M.; visualization, R.A.-A., H.F., T.B.-B., S.S., I.Q. and P.A.-M.; supervision, P.A.-M.; funding acquisition, P.A.-M. All authors have read and agreed to the published version of the manuscript.

## Acknowledgments

The authors thank M. L. Navarro and S. Ramon (IDiBE, Universidad Miguel Hernández) for their excellent technical assistance.

