## Supplementary material for "EXPLORING THE EFFECTS OF METABOLISM-DISRUPTING CHEMICALS ON PANCREATIC α-CELL BIOLOGY: A SCREENING TESTING APPROACH"

| 48 h | RZ | NRU | CFDA-AM |
|---|---|---|---|
| **Control** | 100.00 ± 1.72 | 100.00 ± 0.94 | 100.00 ± 1.01 |
| **BPA 100 pM** | 94.13 ± 2.04 | 98.74 ± 1.23 | 96.92 ± 0.84 |
| **BPA 1 nM** | 92.42 ± 2.00 * | 96.31 ± 0.96 | 96.44 ± 1.02 |
| **BPA 10 nM** | 93.27 ± 2.18 | 97.58 ± 0.87 | 96.97 ± 1.35 |
| **BPA 100 nM** | 90.86 ± 2.00 ** | 97.12 ± 1.03 | 94.27 ± 0.87 ** |
| **BPA 1 µM** | 91.56 ± 1.76 * | 95.43 ± 1.19 * | 94.54 ± 1.24 ** |
| **BPA 10 µM** | 96.26 ± 1.90 | 93.80 ± 1.49 *** | 93.70 ± 1.18 *** |
| | **RZ** | **NRU** | **CFDA-AM** |
| **Control** | 100.00 ± 1.15 | 100.00 ± 1.49 | 100.00 ± 0.69 |
| **BPS 100 pM** | 100.85 ± 1.61 | 103.46 ± 1.36 | 98.92 ± 0.61 |
| **BPS 1 nM** | 97.83 ± 1.14 | 103.67 ± 0.96 | 99.32 ± 0.76 |
| **BPS 10 nM** | 100.93 ± 1.36 | 102.61 ± 1.32 | 100.57 ± 0.49 |
| **BPS 100 nM** | 100.22 ± 1.91 | 103.85 ± 1.67 | 99.27 ± 0.94 |
| **BPS 1 µM** | 97.91 ± 2.07 | 103.17 ± 1.12 | 98.23 ± 1.02 |
| **BPS 10 µM** | 89.91 ± 3.72 | 90.34 ± 4.04 | 97.53 ± 2.04 |
| | **RZ** | **NRU** | **CFDA-AM** |
| **Control** | 100.00 ± 0.61 | 100.00 ± 0.94 | 100.00 ± 0.84 |
| **BPF 100 pM** | 96.66 ± 1.03 | 98.54 ± 1.11 | 97.70 ± 0.95 |
| **BPF 1 nM** | 93.63 ± 1.18 *** | 97.57 ± 1.30 | 94.51 ± 0.97 ** |
| **BPF 10 nM** | 91.94 ± 1.09 **** | 96.39 ± 1.02 | 93.84 ± 1.00 **** |
| **BPF 100 nM** | 92.35 ± 1.02 **** | 98.62 ± 1.01 | 92.49 ± 0.82 **** |
| **BPF 1 µM** | 90.57 ± 1.15 **** | 97.90 ± 1.18 | 91.45 ± 1.06 **** |
| **BPF 10 µM** | 86.61 ± 1.27 **** | 95.01 ± 1.03 * | 91.27 ± 1.15 **** |
| | **RZ** | **NRU** | **CFDA-AM** |
| **Control** | 100.00 ± 0.87 | 100.00 ± 0.75 | 100.00 ± 0.91 |
| **DEHP 100 pM** | 96.88 ± 0.84 | 99.16 ± 0.46 | 98.62 ± 0.79 |
| **DEHP 1 nM** | 93.80 ± 1.27 ** | 99.69 ± 0.88 | 96.72 ± 0.97 |
| **DEHP 10 nM** | 94.70 ± 1.30 * | 99.53 ± 1.40 | 96.16 ± 0.97 * |
| **DEHP 100 nM** | 93.77 ± 1.30 ** | 99.08 ± 0.68 | 96.01 ± 1.02 * |
| **DEHP 1 µM** | 93.25 ± 1.42 ** | 100.09 ± 1.19 | 94.73 ± 0.94 *** |
| **DEHP 10 µM** | 106.54 ± 1.83 ** | 97.31 ± 0.76 | 95.02 ± 0.94 ** |
| | **RZ** | **NRU** | **CFDA-AM** |
| **Control** | 100.00 ± 0.85 | 100.00 ± 1.33 | 100.00 ± 4.02 |
| **PFOS 100 pM** | 96.39 ± 1.33 | 98.99 ± 1.14 | 97.94 ± 1.15 |
| **PFOS 1 nM** | 91.99 ± 1.84 ** | 97.62 ± 1.01 | 96.36 ± 1.73 |
| **PFOS 10 nM** | 85.62 ± 2.37 **** | 88.47 ± 3.45 **** | 93.34 ± 1.92 * |
| **PFOS 100 nM** | 91.97 ± 1.56 ** | 97.37 ± 0.63 | 93.56 ± 2.08 ** |
| **PFOS 1 µM** | 89.13 ± 1.34 **** | 97.42 ± 1.49 | 93.26 ± 0.61 ** |
| **PFOS 10 µM** | 93.94 ± 1.63 * | 97.20 ± 1.84 | 91.74 ± 1.07 *** |
| | **RZ** | **NRU** | **CFDA-AM** |
| **Control** | 100.00 ± 1.29 | 100.00 ± 0.92 | 100.00 ± 0.88 |
| **$CdCl_2$ 100 pM** | 98.90 ± 1.35 | 99.56 ± 1.25 | 98.67 ± 0.99 |
| **$CdCl_2$ 1 nM** | 98.30± 1.56 | 98.12 ± 1.43 | 98.12 ± 1.19 |
| **$CdCl_2$ 10 nM** | 97.55 ± 1.58 | 97.85 ± 1.22 | 97.17 ± 1.20 |
| **$CdCl_2$ 100 nM** | 97.82 ± 1.67 | 95.41 ± 1.29 * | 98.02 ± 1.19 |
| **$CdCl_2$ 1 µM** | 96.31 ± 1.69 | 92.60 ± 1.73 *** | 96.39 ± 1.37 |
| **$CdCl_2$ 10 µM** | 95.16 ± 1.76 | 94.86 ± 1.09 * | 95.93 ± 1.39 |
| | **RZ** | **NRU** | **CFDA-AM** |
| **Control** | 100.00 ± 0.69 | 100.00 ± 0.52 | 100.00 ± 0.50 |
| **DDE 100 pM** | 96.04 ± 1.16 * | 98.81 ± 1.00 | 97.99 ± 0.94 |
| **DDE 1 nM** | 94.26 ± 1.16 *** | 98.33 ± 0.68 | 96.06 ± 0.44 |
| **DDE 10 nM** | 94.00 ± 1.32 *** | 99.19 ± 0.81 | 94.19 ± 1.14 *** |
| **DDE 100 nM** | 94.96 ± 1.04 ** | 101.68 ± 0.70 | 95.44 ± 1.22 * |
| **DDE 1 µM** | 95.03 ± 1.24 ** | 101.69 ± 0.86 | 94.13 ± 1.20 *** |
| **DDE 10 µM** | 95.87 ± 1.04 * | 103.55 ± 0.74 ** | 90.60 ± 1.19 **** |

**Supplemental Table S1.** Viability of pancreatic αTC1-9 cells treated for 48 h with different BPA, BPS, BPF, DEHP, PFOS, $CdCl_2$, or DDE concentrations (100 pM–10 µM) as evaluated by RZ, NR and CFDA-AM assays. n= at least 3 independent experiments. All data are expressed as mean ± SEM. *vs. Control; *p < 0.05, **p < 0.01, ***p < 0.001 and ****p < 0.0001 by one-way ANOVA followed by Dunnet's post hoc test or Kruskal-Wallis followed by Dunn's post hoc test.

| 72 h | RZ | NRU | CFDA-AM |
|---|---|---|---|
| **Control** | 100.00 ± 1.28 | 100.00 ± 0.64 | 100.00 ± 0.75 |
| **BPA 100 pM** | 98.51 ± 1.53 | 100.09 ± 1.28 | 100.00 ± 0.99 |
| **BPA 1 nM** | 94.97 ± 1.43 | 100.27 ± 1.38 | 98.49 ± 1.11 |
| **BPA 10 nM** | 96.22 ± 1.79 | 98.61 ± 1.34 | 97.86 ± 0.92 |
| **BPA 100 nM** | 97.53 ± 1.61 | 100.68 ± 1.30 | 98.22 ± 0.75 |
| **BPA 1 µM** | 95.76 ± 0.94 | 101.80 ± 1.26 | 99.59 ± 0.94 |
| **BPA 10 µM** | 93.17 ± 1.04 ** | 97.64 ± 1.45 | 97.36 ± 1.43 |
| | **RZ** | **NRU** | **CFDA-AM** |
| **Control** | 100.00 ± 1.07 | 100.00 ± 0.70 | 100.00 ± 0.83 |
| **BPS 100 pM** | 98.49 ± 1.03 | 97.28 ± 0.88 | 99.50 ± 0.80 |
| **BPS 1 nM** | 96.04 ± 1.11 * | 94.84 ± 1.21 ** | 98.46 ± 0.58 |
| **BPS 10 nM** | 95.03 ± 1.61 * | 98.90 ± 1.52 | 98.73 ± 1.07 |
| **BPS 100 nM** | 97.62 ± 0.95 | 99.84 ± 1.70 | 97.76 ± 0.94 |
| **BPS 1 µM** | 97.87 ± 1.84 | 98.54 ± 1.47 | 99.34 ± 1.26 |
| **BPS 10 µM** | 89.82 ± 2.97 * | 96.58 ± 2.50 | 97.28 ± 0.89 |
| | **RZ** | **NRU** | **CFDA-AM** |
| **Control** | 100.00 ± 0.61 | 100.00 ± 0.52 | 100.00 ± 0.54 |
| **BPF 100 pM** | 95.10 ± 0.61 * | 98.55 ± 1.05 | 97.90 ± 0.71 |
| **BPF 1 nM** | 95.16 ± 1.04 * | 98.99 ± 0.70 | 97.29 ± 0.63 * |
| **BPF 10 nM** | 92.76 ± 1.04 **** | 97.12 ± 0.99 | 96.21 ± 1.04 ** |
| **BPF 100 nM** | 93.14 ± 1.23 **** | 97.44 ± 1.04 | 95.95 ± 0.88 ** |
| **BPF 1 µM** | 92.30 ± 1.30 **** | 95.83 ± 1.05 | 95.47 ± 0.85 *** |
| **BPF 10 µM** | 90.83 ± 1.43 **** | 95.24 ± 1.16 | 95.40 ± 0.84 *** |
| | **RZ** | **NRU** | **CFDA-AM** |
| **Control** | 100.00 ± 1.39 | 100.00 ± 0.87 | 100.00 ± 1.16 |
| **DEHP 100 pM** | 95.94 ± 1.67 | 99.99 ± 1.20 | 95.93 ± 1.36 |
| **DEHP 1 nM** | 93.96 ± 1.49 * | 99.03 ± 1.11 | 94.08 ± 1.34 * |
| **DEHP 10 nM** | 91.38 ± 1.07 *** | 99.33 ± 1.37 | 94.65 ± 1.33 * |
| **DEHP 100 nM** | 93.84 ± 1.61 * | 99.07 ± 0.97 | 94.07 ± 1.52 * |
| **DEHP 1 µM** | 94.53 ± 1.45 * | 101.80 ± 1.23 | 92.49 ± 1.39 *** |
| **DEHP 10 µM** | 105.00 ± 1.66 | 97.78 ± 1.14 | 92.78 ± 1.30 ** |
| | **RZ** | **NRU** | **CFDA-AM** |
| **Control** | 100.00 ± 1.75 | 100.00 ± 1.97 | 100.00 ± 1.10 |
| **PFOS 100 pM** | 98.71 ± 1.15 | 95.05 ± 2.08 | 97.57 ± 0.92 |
| **PFOS 1 nM** | 98.32 ± 0.81 | 94.99 ± 2.55 | 97.80 ± 0.64 |
| **PFOS 10 nM** | 82.48 ± 3.25 **** | 82.20 ± 3.66 *** | 90.85 ± 1.83 **** |
| **PFOS 100 nM** | 93.06 ± 2.72 | 92.35 ± 2.94 | 95.84 ± 1.38 * |
| **PFOS 1 µM** | 95.63 ± 1.19 | 88.27 ± 2.60 ** | 96.38 ± 0.83 * |
| **PFOS 10 µM** | 99.78 ± 2.33 | 91.78 ± 2.09 * | 98.87 ± 1.01 |
| | **RZ** | **NRU** | **CFDA-AM** |
| **Control** | 100.00 ± 0.96 | 100.00 ± 2.12 | 100.00 ± 0.68 |
| **$CdCl_2$ 100 pM** | 100.90 ± 1.20 | 101.00 ± 2.30 | 99.63 ± 0.76 |
| **$CdCl_2$ 1 nM** | 99.99 ± 0.97 | 100.90 ± 2.61 | 99.07 ± 0.90 |
| **$CdCl_2$ 10 nM** | 97.89 ± 1.26 | 102.90 ± 2.61 | 98.25 ± 0.86 |
| **$CdCl_2$ 100 nM** | 100.60 ± 1.00 | 102.10 ± 2.35 | 98.62 ± 1.29 |
| **$CdCl_2$ 1 µM** | 98.95 ± 1.04 | 101.20 ± 1.53 | 97.93 ± 1.16 |
| **$CdCl_2$ 10 µM** | 95.61 ± 1.45 * | 99.33 ± 1.56 | 97.10 ± 1.27 |
| | **RZ** | **NRU** | **CFDA-AM** |
| **Control** | 100.00 ± 0.46 | 100.00 ± 0.58 | 100.00 ± 0.50 |
| **DDE 100 pM** | 97.84 ± 0.43 ** | 97.93 ± 0.57 | 98.21 ± 0.59 |
| **DDE 1 nM** | 98.75 ± 0.54 | 99.33 ± 0.62 | 98.91 ± 0.53 |
| **DDE 10 nM** | 98.30 ± 0.56 | 99.43 ± 0.69 | 97.87 ± 0.66 |
| **DDE 100 nM** | 98.10 ± 0.61 ** | 99.74 ± 0.65 | 97.33 ± 0.59 ** |
| **DDE 1 µM** | 97.26 ± 0.44 *** | 99.90 ± 0.61 | 97.21 ± 0.57 ** |
| **DDE 10 µM** | 97.02 ± 0.49 *** | 100.10 ± 0.89 | 96.28 ± 0.59 **** |

**Supplemental Table S2.** Viability of pancreatic αTC1-9 cells treated for 72 h with different BPA, BPS, BPF, DEHP, PFOS, $CdCl_2$, or DDE concentrations (100 pM–10 µM) as evaluated by RZ, NR and CFDA-AM assays. n= at least 3 independent experiments. All data are expressed as mean ± SEM. *vs. Control; *$p < 0.05$, **$p < 0.01$, ***$p < 0.001$ and ****$p < 0.0001$ by one-way ANOVA followed by Dunnet's post hoc test, or Kruskal-Wallis followed by Dunn's post hoc test.

| Gene | Forward (5´→ 3´) | Reverse (5´→ 3´) |
|---|---|---|
| *Gcg* | CACTCACAGGGCACATTCAC | TTTGGCAATGTTGTTCCGGTT |
| *Gck* | TTCAGCTTCTGGCCTCCCACAG | AAAACAGCCAGGTCTGGGCAGC |
| *Glut1* | GTGTCGCTGTTTGTTGTAGAG | CAAAGCCAAAGATGGCCACGA |
| *Arx* | GGCCGGAGTGCAAGAGTAAAT | TGCATGGCTTTTTCCTGGTCA |
| *MafB* | ACCAAGGACGAGGTGATCC | CAGGTGATGTTTCTGCTGGA |
| *Foxo1* | AAGAGCGTGCCCTACTTCAA | CTCTTGCCCAGACTGGAGAG |
| *Hprt* | GGTTAAGCAGTACAGCCCCA | TCCAACACTTCGAGAGGTCC |
| *Actb* | GGCTGTATTCCCCTCCATCG | CCAGTTGGTAACAATGCCATGT |
| *Gapdh* | ACACTGAGCAAGAGAGGCCCTA | GGGTGCAGCGAACTTTATTGATGGTATT |

**Supplemental Table S3.** Primer sequences used in RT-qPCR for the study of pancreatic α-cell gene expression.
